# Ablation of *mpeg*+ macrophages exacerbates *mfrp*-related hyperopia

**DOI:** 10.1101/2021.05.03.442504

**Authors:** Zachary J. Brandt, Ross F Collery, Joseph C Besharse, Brian A. Link

## Abstract

**PURPOSE:** Proper refractive development of the eye, termed emmetropization, is critical for focused vision and impacted by both genetic determinants and several visual environment factors. Improper emmetropization caused by genetic variants can lead to congenital hyperopia, which is characterized by small eyes and relatively short ocular axial length. To date variants in only four genes have been firmly associated with human hyperopia, one of which is *MFRP*. Zebrafish *mfrp* mutants also have hyperopia and similar to reports in mice, exhibit increased macrophage recruitment to the retina. The goal of this research was to examine the effects of macrophage ablation on emmetropization and *mfrp*-related hyperopia.

**METHODS:** We utilized a chemically inducible, cell-specific ablation system to deplete macrophages in both wild-type and *mfrp* mutant zebrafish. Spectral-domain optical coherence tomography (SD-OCT) was used to measure components of the eye and determine relative refractive state. Histology, immunohistochemistry, and transmission electron microscopy was used to further study the eyes.

**RESULTS:** While macrophage ablation does not cause significant changes to the relative refractive state of wild-type zebrafish, macrophage ablation in *mfrp* mutants significantly exacerbates their hyperopic phenotype.

**CONCLUSIONS:** Genetic inactivation of *mfrp* leads to hyperopia as well as abnormal accumulation of macrophages in the retina. Ablation of the mpeg1-positive macrophage population exacerbates the hyperopia, suggesting that macrophages are recruited in an effort help preserve emmetropization and ameliorate hyperopia.

## METHODS

### Histology and TEM

Eyes were fixed with 1.0% paraformaldehyde, 2.5% glutaraldehyde, 3.0% sucrose, in 0.06 M cacodylate buffer overnight at 4°C. Samples were then washed in cacodylate buffer and post-fixed with 1% osmium tetroxide and dehydrated by series of methanol washes. Larvae were infused with Epon 812 resin (Electron Microscopy Sciences) through two 15 minute acetonitrile washes followed by 1:1 acetonitrile:Epon incubation for 1 hour, and 100% Epon incubation overnight. Finally, larvae were embedded in 100% Epon and hardened at 65°C for 24 hours. 1μm transverse serial sections through the length of the larvae were cut via microtome and stained with toluidine blue for light microscopy. Light microscopy images were taken using a NanoZoomer 2.0-HT (Hamamatsu Photonics K.K.). For TEM analysis 70nm sections were cut, collected on hexagonal grids and stained with uranyl acetate and lead citrate, followed by imaging on a Hitachi H-600 electron microscope.

### Paraffin Histology

Eyes utilized for paraffin histology were immersed in 4% paraformaldehyde overnight at 4°C and embedded in paraffin blocks for sectioning. 4μm sections were obtained and stained with hematoxylin and eosin for analysis, with serial unstained sections used for immunofluorescent staining. Unstained sections underwent de-paraffinization with xylenes and an ethanol gradient prior to heated antigen retrieval in antigen retrieval solution (Dako). Immunofluorescent staining was then performed as follows.

### Immunofluorescence

Whole-mount immunofluorescent staining was performed on dissected eyecups that were fixed overnight at 4°C in 4% paraformaldehyde. Before whole-mount staining, the lens, cornea, and anterior chamber of the eye was dissected away to allow better access to the tissue. Retinae were washed in PBS to remove fixative. Standard immunostaining followed with 1-hour incubation in blocking solution (2% normal goat serum, 1% TritonX-100, 1% Tween-20 in PBS). Larvae were incubated in primary antibodies overnight in blocking solution at room temperature or 4°C. Embryos were then washed three times for 1 hour in 1% Tween-20 in PBS. Antibody detection was performed using AlexaFluor (488 and 568) conjugated secondary antibodies from Invitrogen at 1:800 dilution in blocking solution overnight at 4°C followed by washes with 1% Tween-20 in PBS.

The following primary antibodies and concentrations were utilized:

1:200 mouse anti-4C4 (Gift from Peter Hitchcock, University of Michigan)
1:500 rabbit anti-Lcp1(L-plastin) (GTX124420, Genetex)
1:500 mouse anti-GFP (Takara, JL-8)
1:500 rabbit anti-GFP (G10362, ABfinity)

### Spectral Domain – Optical Coherence Tomography (SD-OCT)

Zebrafish eyes were imaged using a Bioptigen Envisu R2200 SD-OCT imaging system with a 12 mm telecentric lens (Bioptigen, Morrisville, NC) using a Superlum Broadlighter T870 light source centered at 878.4 nm with a 186.3 nm band width (Superlum, Cork, Ireland). Axial length, lens diameter and retinal radius were measured for populations of zebrafish at 56dpf as previously described ^1^. Both eyes were measured for each fish. In statistical analysis, only the right eye was utilized.

### Eye and body length measurement

Zebrafish eye dimensions were measured as follows: axial length – front of cornea to back of RPE; lens diameter – anterior surface of lens to posterior surface; retinal radius – center of lens to the back of the RPE. Body length was measured from the tip of the head to the end of the trunk (before the caudal fin). Relative refractive error was calculated as previously described^1^.

### Metronidazole (MTZ) treatment

All fish used for macrophage ablation experiments were raised under normal conditions from 0-14dpf. On 14dpf fish were randomly separated into control and experimental groups. Experimental groups were reared in stationary tanks with 7mM MTZ dissolved in fish facility water from 6:00pm to 9:00am daily. From 9:00am – 6:00pm fish were returned to the circulating facility water in 3L tanks for rearing and feeding. From 14dpf – 21dpf fish in the experimental group were kept in 500mL of MTZ for treatments, while 21dpf and older fish were kept in 1L of MTZ for treatments. Untreated control groups are also moved to stationary tanks with equivalent volumes of water during MTZ treatments.

### Zebrafish

All transgenic and mutant lines were generated and maintained in the ZDR genetic background. Wild-type siblings or cousins were used as control groups. All animal experiments were approved by the Institutional Animal Care and Use Committee of the Medical College of Wisconsin.

### Transgenic Lines and Mutant Lines

Tg(*mpeg1*:NTR-eyfp)^2,3^

*Mfrp MW78*^4^

## INTRODUCTION

Emmetropization is the precise regulation of size, morphology, and relative proportions of ocular tissues and is critical for proper refraction of light, and thus clear vision. Improper emmetropization results in either myopia or hyperopia. Myopia is caused by a relative elongation of the axial length and is the more common and better studied refractive error. Hyperopia most often occurs when the axial length is too short for the eye’s focusing apparatus, resulting in light culminating behind the retina. Comparatively less is known regarding the mechanisms of hyperopia.

Variants in just a few genes have been associated with hyperopia. Homozygous or compound heterozygous mutations of *MFRP*, which encodes membrane-type frizzled-related protein, are associated with microphthalmia, high hyperopia, foveoschisis, areas of retinal pigmented epithelium (RPE) atrophy, and optic disc drusen in humans^5–13^. Evidence for the conservation of *MFRP* function comes from multiple animal models. First, two mouse models of spontaneous retinal degeneration, *rd6* and *rdx*, have identified mutations in *Mfrp* as their cause^14,15^. While these mutations resulted in retinal degeneration, initial analysis of homozygous mutants did not find hyperopia. However, examination of the *rd6* model using non-invasive imaging found that these mice indeed have slight decreases in axial length, and that this effect could be rescued by gene therapy^16,17^. Our lab utilized zebrafish to investigate the effects of *mfrp* mutation. In contrast to mice, zebrafish homozygous for *mfrp* mutations do not develop retinal degeneration, but do recapitulate the pronounced hyperopia seen in humans homozygous for *MFRP* alterations^4^. These findings illustrate the role of Mfrp in proper emmetropization and its functional conservation across multiple species.

While both mouse and zebrafish models display hyperopic phenotypes in the absence of Mfrp, neither model fully recapitulates the full spectrum of human *MFRP*-related phenotypes, as mice appear to exhibit only small changes in eye size, failing to develop high hyperopia, and zebrafish mutants do not present photoreceptor degeneration. Intriguingly disruption of *mfrp* in both zebrafish and mice causes accumulation of subretinal macrophages^4,14,18^. Accumulation of retinal macrophages may also occur in human *MFRP-related* pathology. The presence of round yellow-white flecks has been documented in patients with *MFRP*-associated microphthalmia^19^. Under fundus microscopy the subretinal macrophages present in the *Mfrp* mouse models also share this white fleck appearance^14^. These observations suggest that accumulation of retinal macrophages is a unified feature of Mfrp mutations across species.

Based on the accumulation of macrophages seen in two distinct animal models of MFRP-related hyperopia, we hypothesized that retinal macrophages or macrophages within the eye in general, may function to regulate emmetropization. In order to test this hypothesis, we utilized established cell-specific ablation techniques in the zebrafish model. We found that while ablation of macrophages in wild-type fish does not affect basal emmetropization, deletion of this population exacerbates the hyperopia observed in *mfrp* mutant fish. We also investigated changes in proliferation, cell death, and scleral collagen fiber morphology of *mfrp* mutant zebrafish.

## RESULTS

To address the role of macrophages in emmetropization, we sought to deplete the macrophage population during ocular growth and assess potential changes in axial length and refractive error. We chose zebrafish as our model organism as they display a larger shift in refractive state upon *mfrp* deletion than the *mfrp* mutant mouse models, allowing for easier detection of phenotypic change^4,17^. For efficient macrophage ablation we used an established chemical-genetic system^20^. Using macrophage promoter *mpeg1*^2^ we expressed bacterial nitroreductase (NTR) fused to yellow fluorescent protein (eYFP) specifically in macrophages. Fish carrying this transgene are termed *mpeg1:*NTR-eYFP+^3^. On its own, NTR expression does not harm cells; however, it converts the non-toxic prodrug metronidazole (MTZ) into a cytotoxic metabolite that results in autonomous DNA crosslinking and subsequent cell death. This system has been effectively utilized for the ablation of multiple cell types in zebrafish^20^.

### Ablation of macrophages in WT zebrafish does not significantly impact emmetropization

To examine the effects of *mpeg1*+ cell ablation on emmetropization we allowed *mpeg1:*NTR-eYFP+ fish and their non-transgenic wild-type cousins to grow to 8 weeks of age, or more precisely 56 days post fertilization (dpf), with or without MTZ treatment. MTZ was administered through daily bath application starting at 14dpf. We chose to start treatment after 14dpf to allow for the normal development and organization of the retina, and to focus solely on ocular growth as it pertains to eye size and refractive state. Efficient ablation of macrophages in transgenic fish was confirmed by imaging eYFP expressing cells in retinal flat-mount preparations, resulting in near complete depletion (Fig. S1A-A”). When fish reached 56dpf they were anesthetized and imaged via spectral domain optical coherence tomography (SD-OCT) to measure axial length, lens diameter, and focal length (Fig. 1A), which are used to calculate refractive error as previously described^1^.

**Figure 1.**
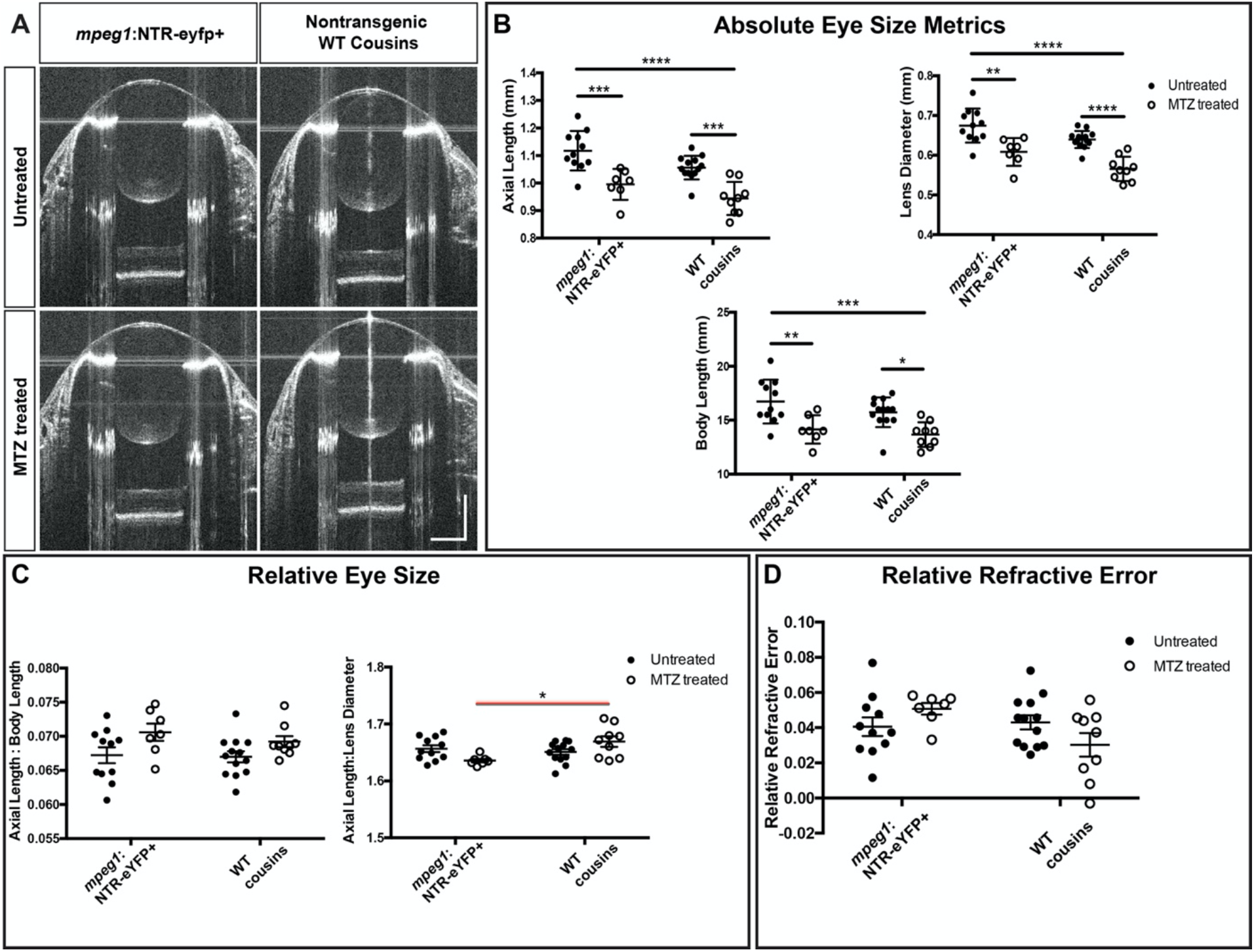
Ablation of macrophages in WT zebrafish does not significantly impact emmetropization. **(A)** Representative SD-OCT B-scans from the center of *mpeg1:*NTR-eYFP+ eyes and their WT cousins with or without MTZ treatment. **(B)** Axial length, Body length, and Lens diameter of *mpeg1:*NTR-eYFP+ fish and their WT cousins with or without MTZ treatment. **(C)** Axial length normalized to Body length and Lens Diameter. **(D)** Relative Refractive error. 2-way ANOVA was used for statistical analysis for B-D with p-values shown from Tukey’s multiple comparisons for *post hoc* analysis. Red bars indicate statistical significance likely due to *mpeg1*+ cell ablation. * = p<0.05, ** = p<0.01, *** = p<0.001, **** = p<0.0001.

MTZ treatment resulted in overall smaller zebrafish, as seen by decreases to axial length, lens diameter, and body length (Fig. 1B). This was true in both the presence and absence of *mpeg1:*NTR-eYFP expression, suggesting that MTZ treatment causes developmental delay or slower overall body growth independent of cell-specific ablation. Because of this nonspecific effect we standardized axial length measurements to body length, as well as lens diameter (Fig. 1C). We have previously shown that both of these measurements correlate linearly with axial length in wild-type fish and can be used for normalization for fish of different sizes^1^. Statistical analysis by 2-way ANOVA indicated a significant effect of MTZ treatment on axial length normalized to body length, but no significant interaction with *mpeg1:*NTR-eYFP+ expression; *post hoc* analysis revealed there were not significant differences between any groups (Fig. 1C). In contrast, 2-way ANOVA of axial length normalized to lens diameter found a significant interaction between presence of the *mpeg1:*NTR-eYFP transgene and MTZ treatment suggesting a possible effect of macrophage ablation on relative eye size. However, the relative eye size of *mpeg1:*NTR-eYFP+ fish treated with MTZ was not statistically different from their *mpeg1:*NTR-eYFP+ untreated siblings (Fig. 1C). To more specifically assess the refractive state of these eyes, we calculated refractive error as previously described^1^. Again statistical analysis by 2-way ANOVA found a significant interaction between presence of the *mpeg1:*NTR-eYFP transgene and MTZ treatment. This suggested that macrophage ablation can affect refractive error, but again *post hoc* multiple comparisons did not reveal significant differences between groups (Fig. 1D). Taken together these results suggest that *mpeg*+ macrophage ablation does not significantly alter emmetropization in wild-type zebrafish.

### Macrophage ablation in *mfrp* mutants

While macrophage ablation did not significantly alter the relative refractive error of WT zebrafish, the previously documented macrophage accumulation in *mfrp* mutants led us to hypothesize that they may still play a role in this pathologic hyperopic state. To evaluate this, we employed the same macrophage ablation strategy on *mfrp* mutant and heterozygous siblings. Again, highly efficient ablation of wild-type macrophages was observed by *mpeg1:*NTR-eYFP expression, as well as using the 4C4 antibody termed 4C4 which labels macrophages and microglial cells in zebrafish (Fig. 2A-B”’, E, F)^21^. While nearly all transgene expressing cells are depleted, some 4C4+ cells escape ablation likely due to incomplete overlap in expression of the 4C4 antigen and *mpeg1* transgene. As expected, *mfrp* mutant fish showed significant increases in *mpeg1:*NTR-eYFP+ cells, with focal increases at the central retina (Fig. 2C-C’”). In *mfrp*-/- fish we saw depletion of *mpeg1:*NTR-eYFP expressing cells; however, the effect was not as complete as in wild-type fish (Fig. 2C, D, E). 4C4 cell counts were not significantly decreased (Fig. 2C’, D’, F). While the ablation of our *mpeg1:*NTR-eYFP expressing cells remained efficient and significant, these results reveal that some macrophages which accumulate in *mfrp*-/- eyes remain.

**Figure 2.**
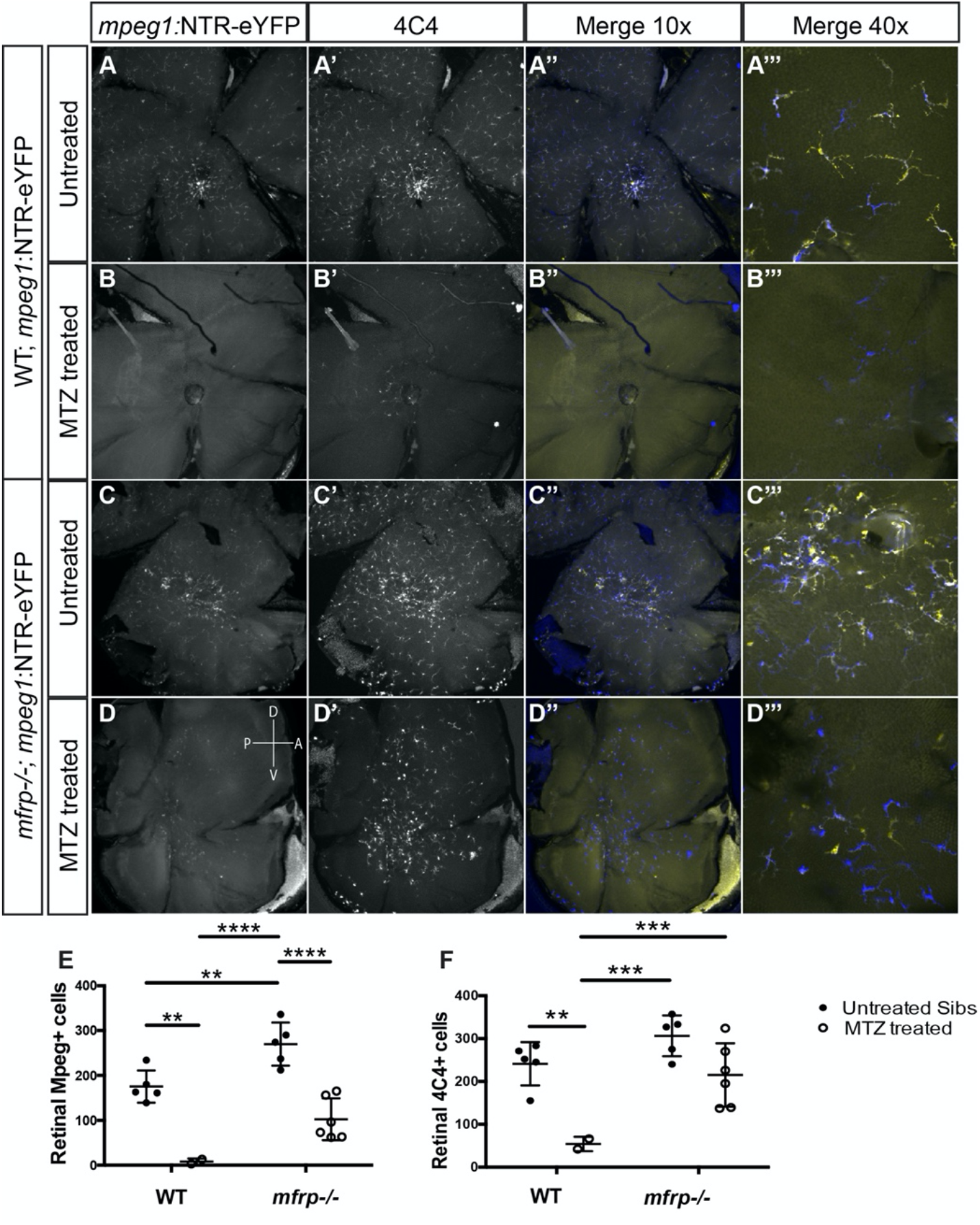
Efficient macrophage ablation in the *mfrp* retina. **(A-D)** *mpeg1:*NTR-eYFP expression in WT and *mfrp*-/- retina flat-mount preparations with and without MTZ treatment. **(A’-D’)** Immune staining for 4C4 antibody marking macrophages. **(A”-D”)** Merged and colorized images of A-D’ with blue representing 4C4 and yellow representing YFP. **(A’”-D’”)** Higher magnification images of A”-D”. **(E, F)** Cell counts of *mpeg1:*NTR-eYFP+ cells (E) or 4C4+ cells (F) in WT and *mfrp*-/- retina flat-mount preparations with and without MTZ treatment. 2-way ANOVA was used for statistical analysis. Sidak’s multiple comparisons for post hoc analysis. * = p<0.05, ** = p<0.01, *** = p<0.001, **** = p<0.0001.

To obtain more detailed spatial information regarding the macrophage accumulation in *mfrp* mutants, as well as their ablation, we performed immunofluorescent staining on histological sections. We labeled eYFP+ cells using a GFP antibody and co-stained for lymphocyte cytosolic protein 1 (Lcp1), as an additional macrophage marker ^22^. Nuclei were stained with 4’,6-diamidino-2-phenylindole (DAPI) to identify the retinal layers. In *mfrp+/-; mpeg1:*NTR-eyfp+ untreated control fish, eYFP+ cells are seen sporadically across all layers of the retina. The ganglion cell layer (GCL) and inner nuclear layer (INL) contain the highest frequency of these cells, but eYFP+ cells can also be found in the outer nuclear layer (ONL). In control fish nearly all eYFP+ cells observed are also Lcp1+ (Fig. 3A-B’”, I-K, M-O). The strong endogenous fluorescence of the photoreceptor layer precluded accurate quantitation of eYFP+ cells in the outer layer; however, Lcp1 staining showed small numbers of macrophages present there as well (Fig. 3L). MTZ treatment efficiently reduced or completely depleted eYFP+ cells across all layers of *mfrp+/-; mpeg1*:NTR-eyfp+ eyes (Fig. 3C-D”’, M-O). Though seemingly reduced, sporadic Lcp1+ cells remained at slightly higher levels than eYFP+ cells, revealing that some macrophages escape ablation. In *mfrp-/-; mpeg1:*NTR-eyfp+ untreated eyes, both eYFP+ and Lcp1+ cells were noted across all layers of the retina (Fig. 3E-F’”). However, statistical analysis showed a significant increase in the frequency of both eYFP+ and Lcp1+ cells only in the ONL (Fig. 3K, O). MTZ treatment was similarly effective in *mfrp-/-; mpeg1:*NTR-eyfp+ fish, as eYFP+ cells were greatly reduced or completely depleted across all layers of the retina (Fig. 3M-O). Lcp1+ cells were also decreased in the INL and ONL (Fig. 3J, K). These data highlight the efficient ablation of macrophages across all layers of the eye and identify the ONL as the specific location of increased macrophage presence in *mfrp*-/- fish.

**Figure 3.**
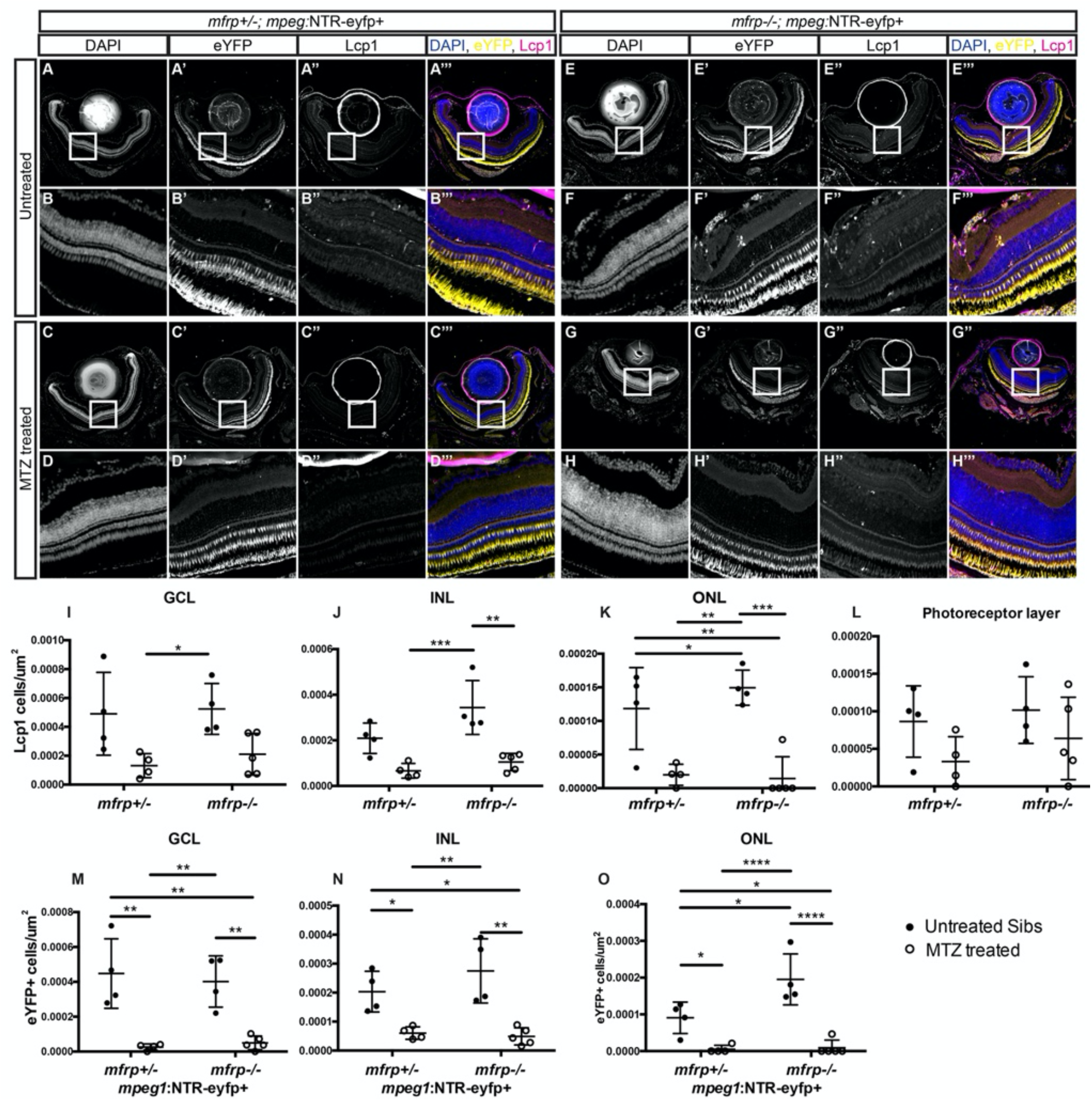
Distribution of macrophage accumulation and ablation across *mfrp+/-;* and *mfrp*-/- retinae. **(A-H)** Representative images of central retina sections from *mfrp+/-; mpeg1:*NTR-eYFP+ and *mfrp-/-; mpeg1*:NTR-eYFP+ MTZ treated and untreated fish. (A-H) Grayscale DAPI images at low (A, C, E, G) and high magnification (B, D, F, H). (A’-H’) Grayscale eYFP images at low (A’, C’, E’, G’) and high magnification (B’, D’, F’, H’). (A”-H”) Grayscale Lcp1 images at low (A”, C”, E”, G”) and high magnification (B”, D”, F”, H”). Colorized merged images with DAPI in blue, eYFP in yellow, and Lcp1 in magenta; images are at low (A”’, C”’, E”’, G”’) and high magnification (B”’, D”’, F”’, H”’). **(I-L)** Quantification of the number of Lcp1+ cells per um^2^ in the ganglion cell layer (I), inner (J) and outer (K) nuclear layer, and photoreceptor layer (L). **(M-O)** Quantification of the number of eYFP+ cells per um^2^ in the ganglion cell layer (M), inner (N) and outer (O) nuclear layer; error bars = standard deviation. 2-way ANOVA was used for statistical analysis for I-O with p-values shown from Tukey’s multiple comparisons for *post hoc* analysis. * = p<0.05, ** = p<0.01, *** = p<0.001, **** = p<0.0001.

### Macrophage ablation exacerbates hyperopia in *mfrp*-/- zebrafish

We again utilized SD-OCT to image and measure the various metrics of *mfrp-/-; mpeg1:*NTR-eYFP+ eyes compared to the eyes of *mfrp+/-; mpeg1:*NTR-eYFP+ siblings with and without MTZ treatment (Fig. 4A). Similar to wild-type fish, MTZ treatment slowed overall growth in both *mfrp*+/-, and *mfrp*-/- fish (Fig. 4B, Body Length). This effect was independent of *mpeg1:*NTR-eYFP expression (Fig S2A-B).

**Figure 4.**
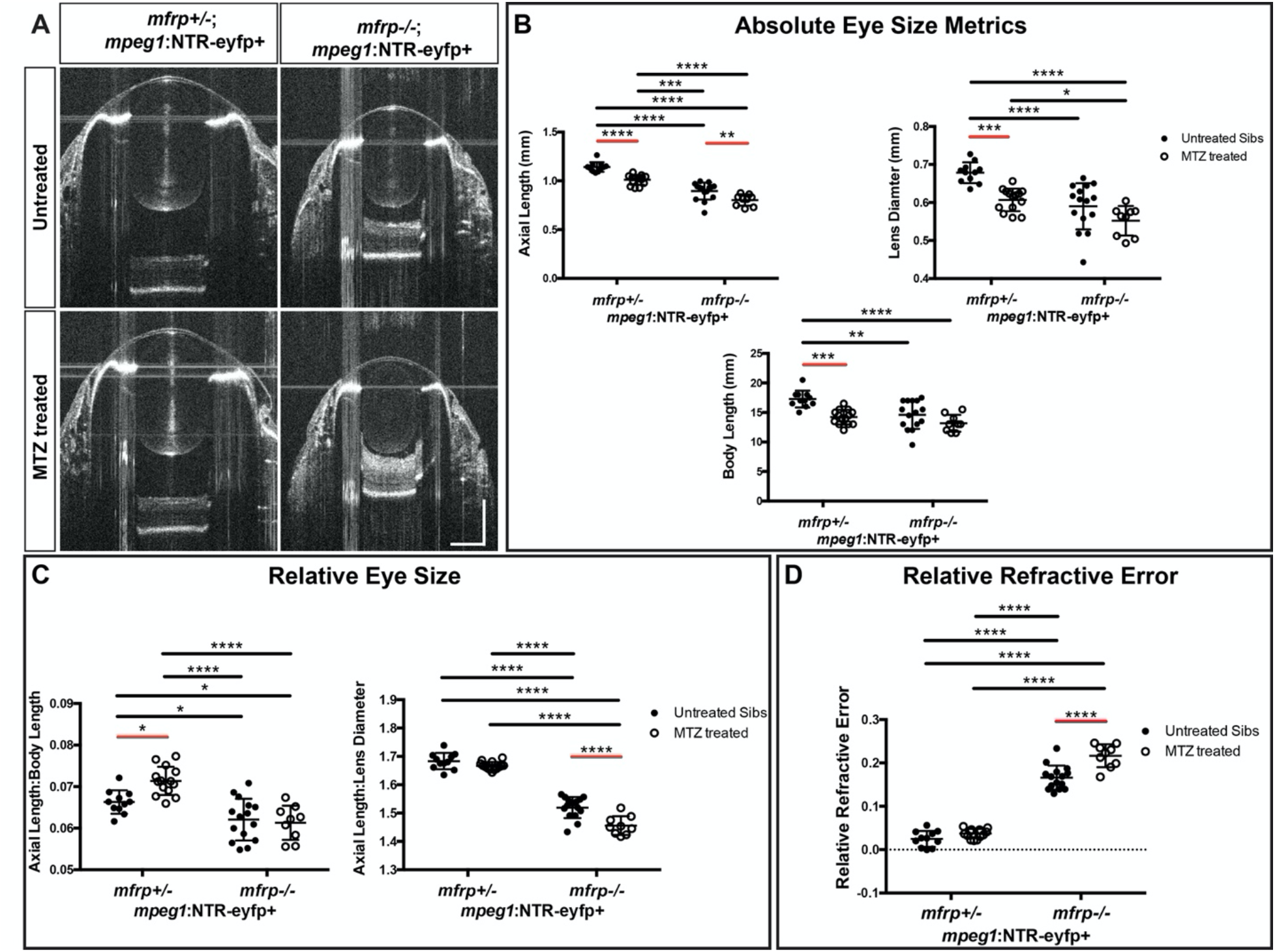
Macrophage ablation exacerbates hyperopia in *mfrp*-/- zebrafish. **(A)** Representative SD-OCT B-scans from the center of mfrp+/-; *mpeg1:*NTR-eYFP+ eyes and their mfrp-/-; *mpeg1:*NTR-eYFP+ siblings with or without MTZ treatment. **(B)** Eye size metrics of *mfrp+/-; mpeg1:*NTR-eYFP+ and *mfrp-/-; mpeg1:*NTR-eYFP+ fish with or without MTZ treatment. **(C)** Axial length normalized to Body length and Lens Diameter. **(D)** Relative Refractive error. 2-way ANOVA was used for statistical analysis with p-values shown from Tukey’s multiple comparisons for *post hoc* analysis. Error bars = standard deviation. Red bars indicate statistical significance due to MTZ treatment. * = p<0.05, ** = p<0.01, *** = p<0.001, **** = p<0.0001.

As expected, *mfrp*-/- fish had significantly reduced relative eye size compared to their *mfrp*+/- siblings, when normalized to either body length or lens diameter as determined by 2-way ANOVA (p<0.0001) (Fig. 4C). Relative refractive error was also significantly increased in *mfrp*-/- fish compared to *mfrp*+/- siblings as previously reported (Fig. 4D)^4^. These results confirm that loss of *mfrp* leads to hyperopia in zebrafish and reveal that *mpeg*+ cell ablation does not rescue this phenotype, but instead appears to exacerbate the hyperopia.

To normalize for the overall growth effect of MTZ treatment and determine relative ocular metrics, we again standardized axial length to both body length and lens diameter. When assessing the effect of MTZ treatment on relative eye size, we found that MTZ treatment affected the axial length to body length ratio of non-transgene control animals (p=0.0043, 2-way ANOVA) (Fig. S2C). While this might suggest a potential nonspecific effect of MTZ treatment on relative eye size, differences were not observed for axial length relative to lens diameter in these non-transgenic control fish (p=0.3664, 2-way ANOVA) (Fig. S2C). Given that body length does not impact emmetropization, while lens diameter directly impacts this process, we concluded that MTZ treatment alone does not alter emmetropization in a nonspecific fashion.

When we assessed the effect of macrophage ablation on the relative eye size of *mfrp+/-; mpeg1:*NTR-eYFP+ compared to *mfrp-/-; mpeg1:*NTR-eYFP+ eyes we found that MTZ treatment affected relative eye size (Fig. 4C; axial length to lens diameter; p<0.0001, 2-way ANOVA). 2-way ANOVA of axial length to lens diameter ratios also revealed a significant interaction between genotype and MTZ treatment (p=0.0158). *post-hoc* analysis confirmed that *mfrp-/-; mpeg1:*NTR-eYFP+ fish had a significantly decreased axial length to lens diameter ratio compared to their untreated *mfrp-/-; mpeg1:*NTR-eYFP+ siblings (Fig. 4C). Again, nonspecific effects were not observed for axial length relative to lens diameter in animals that lacked the *mpeg1:*NTR-eYFP transgene (p=0.3664, 2-way ANOVA) (Fig. S2C), suggesting that the change in relative eye size in *mfrp*-/- fish was specific to *mpeg1*+ cell ablation.

Finally, relative refractive error was also significantly affected by MTZ treatment in *mpeg:*NTR-eYFP expressing fish (Fig. 4D; p<0.0001, 2-way ANOVA), but not in non-transgenic controls (p=0.2646, 2-way ANOVA). Similar to the changes in axial length relative to lens diameter, 2-way ANOVA also revealed a significant interaction between genotype and MTZ treatment (p=0.0043). *post-hoc* analysis showed that the reduction in axial length to lens diameter ratio translated to a significant increase in relative refractive error in *mfrp-/-; mpeg1:*NTR-eYFP+ fish compared to their untreated *mfrp*-/- siblings (Fig. 4D). These results demonstrate that macrophage ablation significantly exacerbates the hyperopic phenotype of *mfrp* mutant zebrafish, indicating that the accumulation of macrophages seen in *mfrp*-related hyperopia may be recruited to ameliorate the microphthalmia.

### Macrophage ablation does not significantly alter retina morphology

As *mpeg1*+ cell ablation affected relative eye size and refractive error we sought to gain possible mechanistic insight into these changes by assessing overall eye and retinal morphology. We assessed the effects of macrophage ablation on overall retina morphology by histology. Hematoxylin and eosin-stained paraffin sections revealed normal retinal morphology following macrophage ablation (Fig. 5A-B’). We did note that MTZ treated eyes displayed more uniform and dispersed melanin throughout the cells of the RPE (Fig. 5A’, B’). The rod outer segment layer also appears shorter when compared with untreated siblings, possibly due to contraction of the myoid. These morphological changes are reminiscent of the retinomotor changes seen in dark adapted zebrafish.

**Figure 5.**
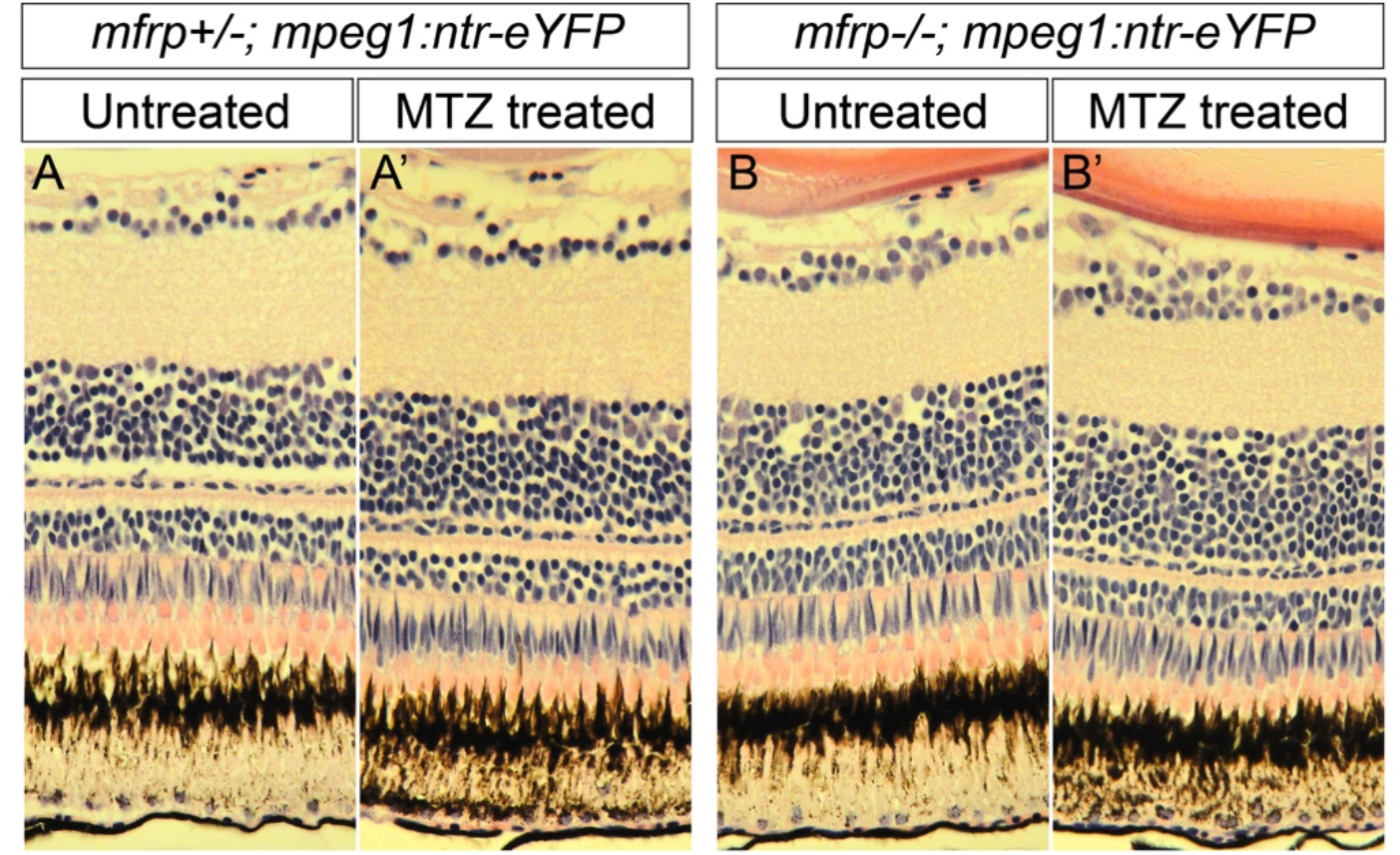
Macrophage ablation does not significantly alter retina morphology. **(A-B’)** H&E stained paraffin histology of the central retina in *mfrp+/-; mpeg1:*NTR-eYFP+ (A-A’), and *mfrp-/-; mpeg1:*NTR-eYFP+ (B-B’) with and without MTZ treatment.

### Neither proliferation nor cell death underlie exacerbated *mfrp*-related hyperopia after macrophage ablation

We hypothesized that the exacerbated changes in relative eye size seen in *mfrp* mutants could be due to altered cell proliferation and/or death. To assess changes in proliferation we performed immunohistochemistry for proliferative cell nuclear antigen (PCNA), along with DAPI to label nuclei. In *mfrp+/-; mpeg1*:NTR-eYFP+ eyes, PCNA+ nuclei are found frequently within both the INL and ONL, as well as the ciliary marginal zone. In comparison, *mfrp-/-; mpeg1:*NTR-eYFP+ fish have similar PCNA+ cell frequency within the INL to their heterozygous siblings, but significantly decreased frequency of PCNA+ nuclei in the ONL (Fig. 6A-B”, E-F”, I-J). No changes in the ciliary marginal zone were noted under any condition. MTZ treatment did significantly decrease the percentage of PCNA+ nuclei in the ONL of *mfrp+/-; mpeg1:*NTR-eYFP+ fish compared to their untreated siblings (Fig. 6C-D”, J). PCNA analysis also revealed that *mfrp*-/- had reduced number of proliferative cells in the ONL, although, MTZ treatment on *mfrp-/-; mpeg1:*NTR-eYFP+ animals did not exacerbate this effect (Fig. 6G-H”, J).

**Figure 6.**
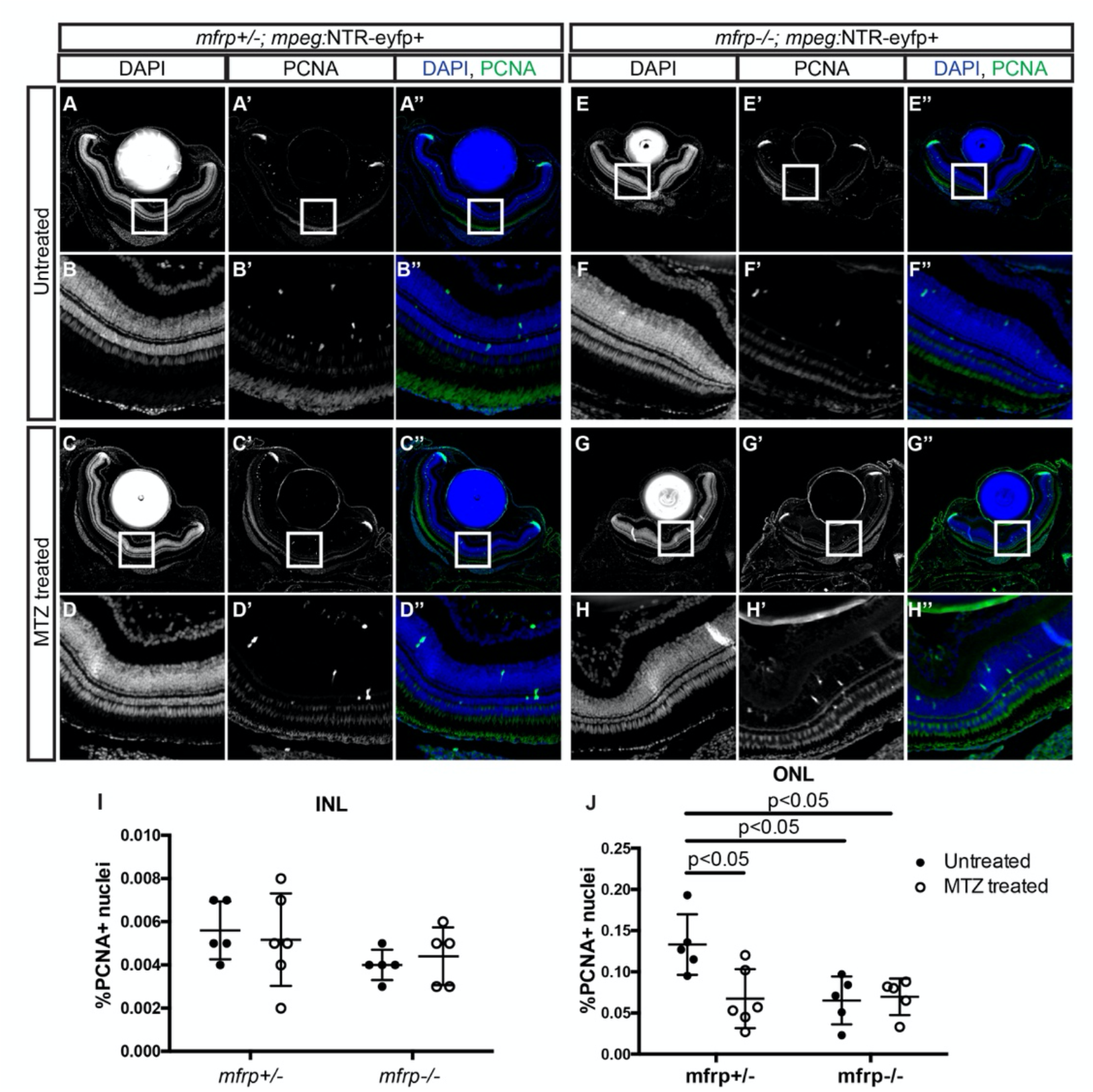
Proliferative effects of *mfrp* deletion and macrophage ablation. **(A-H)** Representative images of central retina sections from *mfrp+/-; mpeg1:*NTR-eYFP+ and *mfrp-/-; mpeg1:*NTR-eYFP+ MTZ treated and untreated fish. (A-H) Grayscale DAPI images at low (A, C, E, G) and high magnification (B, D, F, H). (A’-H’) Grayscale PCNA images at low (A’, C’, E’, G’) and high magnification (B’, D’, F’, H’). (A”-H”) Colorized merged images with DAPI in blue and PCNA in green; images are at low (A”, C”, E”, G”) and high magnification (B”, D”, F”, H”). **(I-J)** Quantification of the percentage of PCNA+ nuclei in the inner (I) and outer (J) nuclear layer; error bars = standard deviation. 2-way ANOVA was used for statistical analysis for I, J with p-values shown from Tukey’s multiple comparisons for *post hoc* analysis.

Alternative to changes cell proliferation, cell death caused by mpeg+ cell ablation could lead to the decrease in relative eye size observed in *mfrp* mutants. To assess cell death, we performed terminal deoxynucleotidyl transferase dUTP nick end labeling (TUNEL). We found little to no TUNEL positive cells regardless of genotype or treatment condition, suggesting that apoptosis was not causing the altered phenotype in *mfrp* mutant eyes (Fig. S3A-C’”). Consistent with this result, DAPI staining did not reveal pyknotic nuclei for any genotype.

These results demonstrate that while macrophage ablation may affect a small number of proliferative cells in the retina it does not affect apoptosis, and neither cellular process appears to underlie the exacerbated hyperopia measured in macrophage ablated *mfrp*-/- fish.

### Macrophage ablation alters collagen bundle size

Past research suggests that scleral collagen synthesis, degradation, and crosslinking play important roles during emmetropization, and this balance is altered during myopia ^23,24^. We hypothesized that collagen fibers in the sclera of *mfrp* mutants may be altered and contribute to their improper emmetropization. To test this, we imaged collagen fibers in the central posterior sclera by transmission electron microscopy (TEM). By measuring and plotting the frequency distribution of collagen fiber diameters in the posterior sclera we see that the posterior scleral tissue in untreated *mfrp-/-; mpeg1:*NTR-eYFP+ fish appears to contain a slightly wider distribution of collagen bundle sizes with a small increase in the proportion of collagen fibers with larger diameters compared to their untreated *mfrp+/-; mpeg1:*NTR-eYFP+ siblings (Fig. 7A-B). MTZ treatment exacerbates these differences (Fig. 7C-D). When compared with their untreated siblings, MTZ treated *mfrp+/-; mpeg1:*NTR-eYFP+ fish display a clear shift in collagen bundle diameter increasing the frequency of smaller collagen bundles and decreasing the frequency of larger bundles (Fig. 7E). These results suggest that macrophage ablation can affect scleral collagen bundle size. Whether or not these changes in scleral collagens underlie the exacerbated *mfrp*-related hyperopia upon macrophage ablation is unclear.

**Figure 7.**
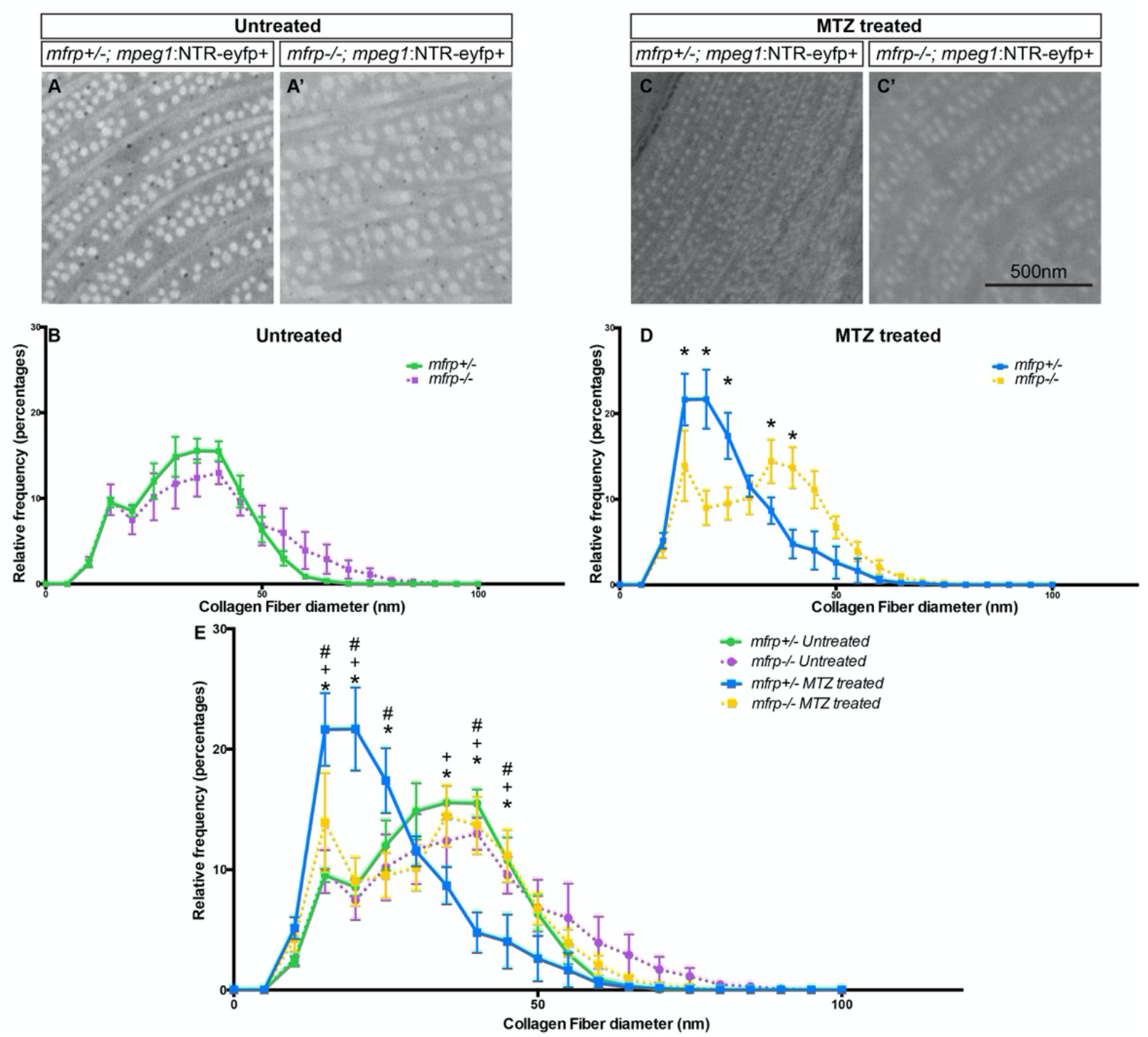
Collagen Fibril Diameter in *mfrp*+/- and *mfrp*-/- sclera with and without macrophage ablation. **(A)** Representative examples of collagen fibrils in the central posterior sclera of *mfrp+/-; mpeg1:*NTR-eYFP+ (A) and *mfrp-/-; mpeg1:*NTR-eYFP+ (A’) untreated fish. **(B)** Frequency distribution of collagen fiber diameter with y-axis = relative frequency as a percentage and x-axis = collagen fiber diameter in 5nm bins. *mfrp*+/- = green line, *mfrp*-/- = purple line. *mfrp*+/- n=3; *mfrp*-/- n=4. **(C)** Representative examples of collagen fibrils in the central posterior sclera of *mfrp+/-; mpeg1:*NTR-eYFP+ (C) and *mfrp-/-; mpeg1*:NTR-eYFP+ (C’) MTZ treated fish. **(D)** Frequency distribution of collagen fiber diameter with y-axis = relative frequency as a percentage and x-axis = collagen fiber diameter in 5nm bins. *mfrp*+/- = blue line, *mfrp*-/- = yellow line. *mfrp*+/- n=5; *mfrp*-/- n=5. Multiple t-tests used for statistical analysis. * denotes significance of p<0.001. **(E)** Combined graph of (B) and (D). Errors bars = standard deviation throughout figure. 2-Way ANOVA with Tukey Multiple comparisons used for statistical analysis. * denotes significant difference between *mfrp*+/- MTZ treated and *mfrp*-/- MTZ treated, + denotes significant difference between *mfrp*+/- MTZ treated and *mfrp*+/- Untreated, and # denotes significant difference between *mfrp*+/- MTZ treated and *mfrp*-/- Untreated. p<0.05 for all significant differences in E.

## DISCUSSION

We find that depletion of macrophages exacerbates the *mfrp*-related hyperopia in zebrafish. Additionally, we observe that while macrophages affect the development of *mfrp*-related hyperopia, the absence of the mpeg-1 population does not significantly alter wild-type emmetropization. We go on to investigate further how macrophage ablation might affect morphology and growth of retina and sclera, examining changes in retinal cell proliferation as well as changes to scleral collagen fibers.

We noted the nonspecific effects of MTZ treatment on the overall size of the fish and accounted for these changes in our analysis. MTZ could cause additional non-obvious effects. MTZ itself is an antibiotic that was originally used for the treatment of trichomoniasis^25,26^, and continues to be used for combating anaerobic infections^27^. Daily antibiotic washes on zebrafish likely have a profound effect on the microbiome of the fish and could result in unintended effects. Metronidazole has also been shown to effect circadian rhythm increasing the expression of core clock genes in the skeletal muscle of germ-free mice^28^. However, this MTZ based ablation technique has been used to study circadian rhythms of visual sensitivity in zebrafish and no alterations to circadian rhythm were reported in non-transgenic MTZ treated fish^29^. To control for potential nonspecific effects of MTZ, all non-transgenic control groups were treated exactly as transgene expressing experimental groups and underwent the same MTZ treatment regimen.

Our ablation technique is limited to the cells expressing the *mpeg1*:NTR-eYFP transgene. In that context, this broad macrophage population is efficiently ablated by the action of the transgene. Still, 4C4+ macrophages remained following MTZ treatment, suggesting non-mpeg+ macrophages persist. Likewise our histological analysis demonstrated lcp1+ macrophages can also persist, although at reduced numbers. These findings highlight the heterogeneity of the population of monocyte-like cells in the retina, whether macrophage or microglia. It is clear that the *mpeg1:NTR-eYFP* transgene, while marking a broad population of macrophages, does not express in all retinal monocytes. Notably, both mpeg1 positive and negative macrophages were elevated in the *mfrp* mutant retina. Future work into the importance of macrophage sub-types could prove useful for identifying their possible role in emmetropization, and potential additional effects on *mfrp* mutant pathogenesis. It seems likely that either the loss of Mfrp or the hyperopic condition of the eye alters the state of macrophages in the retina, possibly leading to activation or recruitment of a separate sub-population.

Significantly, we found alteration to scleral collagen fibril size in *mfrp* mutants, as well as the modulation of collagen fibril size by macrophage ablation. Recent work suggests that collagen bundle size is dynamically regulated during emmetropization, and these alterations in size can change their mechanical properties^23^. Recently a change in scleral collagen dynamics has been partially attributed to an increased presence of scleral macrophages in a visual form-deprived mouse model of myopia^24^. These investigators report that macrophages appear to be recruited to the sclera of form-deprived myopic mice by increased scleral expression of c-c motif chemokine ligand-2 (Ccl2). The authors suggest that upon recruitment the macrophages are then partially responsible for the secretion of matrix metalloproteinase-2 (Mmp-2) in the sclera, resulting in extracellular matrix remodeling that contributes to the development of myopia. One possible hypothesis that stems from these data is that as the retina grows in size it exerts force on the sclera, and collagen remodeling alters scleral compliance, allowing axial elongation of the eye. In the case of *mfrp* mutant eyes, increased collagen bundle diameter may result in increased tensile stiffness of the sclera and prevent proper axial elongation. Indeed, we observed an upward trend in the percentage of collagen fibrils with larger diameter in *mfrp*-/- sclera when compared to their *mfrp*+/- siblings. In further support of these ideas, retinal folds are observed in *mfrp*-/- eyes^4^. Perhaps the retina itself continues to expand while sclera cannot, and this forces the retina to fold in on itself. This would be of particular relevance to the foveoschisis seen in some patients with *MFRP*-related microphthalmia. Furthermore, MTZ treatment resulted in a clear shift of collagen fibril diameter distribution in *mfrp+/-; mpeg1:NTR-eYFP*+ fish, suggesting that the presence of macrophages can affect collagen fibril size. Supporting a direct role for macrophages, it has been reported that macrophages have the ability to directly secrete collagen in the context of scar formation after cardiac injury in zebrafish^30^. Further work is required to define the precise relationship between collagen fibril size and emmetropia, as it currently remains unclear if this alteration to scleral collagen in *mfrp* mutants is an underlying cause of improper refractive state or simply a response.

The most significant finding in this study is the role of the *mpeg1*+ cell population in pathologic hyperopia, but not for wild-type emmetropization. To date very little is known regarding the function of ocular macrophages in either normal or aberrant emmetropization. Our results therefore mark an initial understanding for the role of macrophages in pathologic hyperopia, and more specifically *mfrp*-related hyperopia in zebrafish.

## ACKNOWLEDGEMENTS

We thank David Parichy (University of Virgina, Charlottesville) for sharing the *mpeg1*:NTR-eyfp transgenic line, Clive Wells (Director of the MCW Electron Microscopy core) for assistance with electron microscopy, and Pat Cliff and Erin Bentley for providing zebrafish husbandry.

---

**Figure S1.**
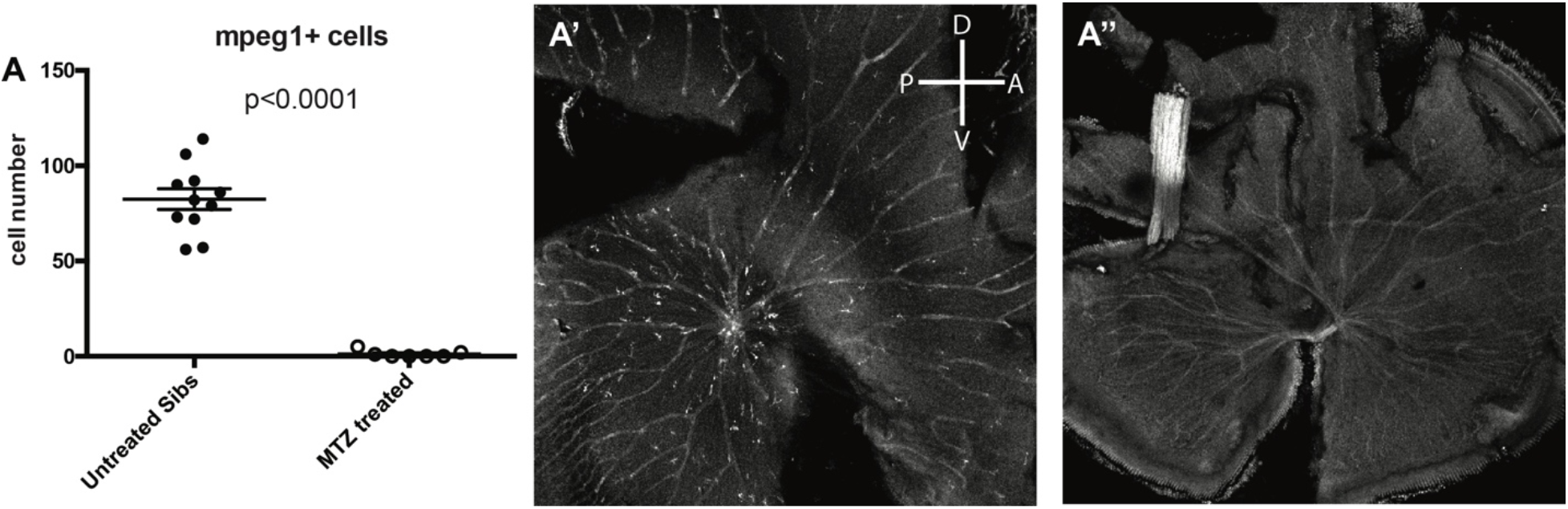
Efficient ablation of macrophages in WT eyes. **(A)** Number of *mpeg1:*NTR-eYFP+ cells identified in the retina with and without MTZ treatment. **(A’,A”)** Representative images of *mpeg1:*NTR-eYFP+ retina flatmounts with(A’) or without(A”) MTZ treatment. Unpaired t-test was used for statistical analysis in A.

**Figure S2.**
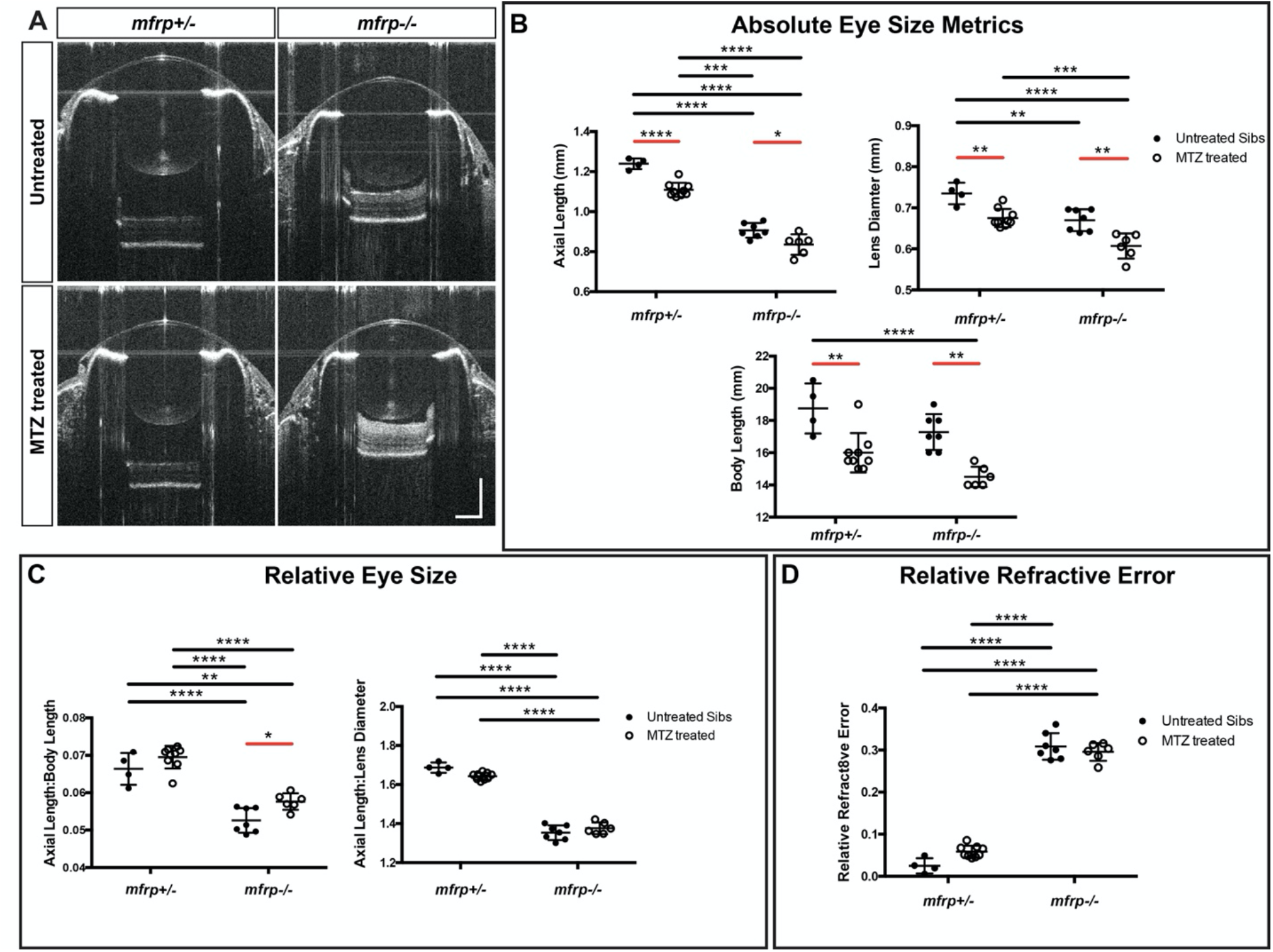
Nonspecific Effects of MTZ treatment on *mfrp* Eye Metrics. **(A)** Representative SD-OCT B-scans from the center of sibling *mfrp*+/- and *mfrp*-/- eyes with or without MTZ treatment. **(B)** Eye size metrics of *mfrp*+/- and *mfrp*-/- fish with or without MTZ treatment. **(C)** Axial length normalized to Body length and Lens diameter. **(D)** Relative Refractive error. 2-way ANOVA was used for statistical analysis with p-values shown from Tukey’s multiple comparisons for *post hoc* analysis. Error bars = standard deviation. Red bars indicate statistical significance due to MTZ treatment. * = p<0.05, ** = p<0.01, *** = p<0.001, **** = p<0.0001.

**Figure S3.**
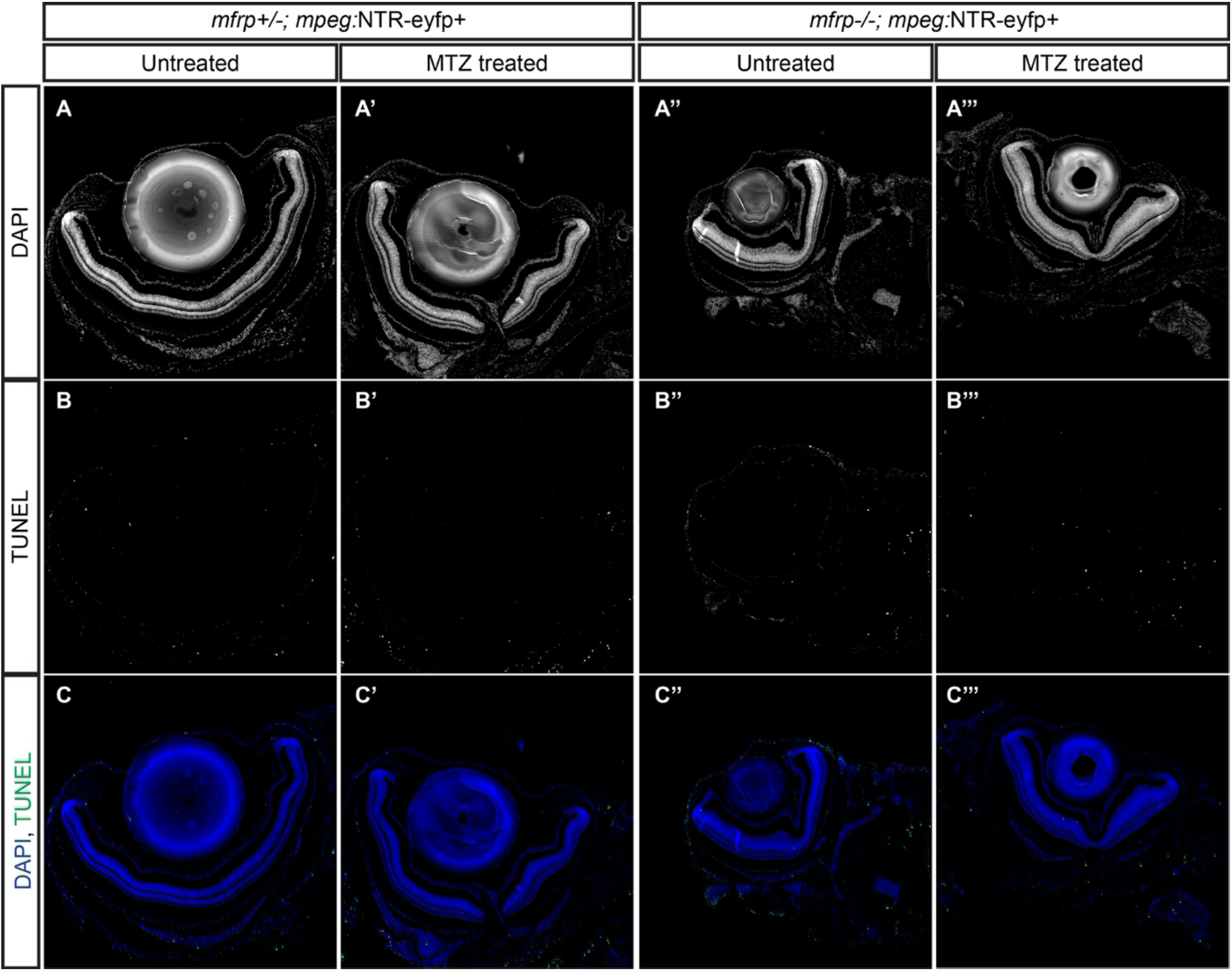
Macrophage ablation causes no appreciable change in retinal cell death. **(A-A”’)** Grayscale DAPI images retina sections from *mfrp +/-; mpeg:*NTR-eyfp+ and *mfrp-/-; mpeg:*NTR-eyfp+ MTZ treated and untreated eyes. **(B-B’”)** Grayscale images of TUNEL stained sections from A. **(C-C’”)** Colorized merged images with DAPI in blue and TUNEL in green.

## REFERENCES

1. Collery, R. F., Veth, K. N., Dubis, A. M., Carroll, J. & Link, B. A. Rapid, Accurate, and Non-Invasive Measurement of Zebrafish Axial Length and Other Eye Dimensions Using SD-OCT Allows Longitudinal Analysis of Myopia and Emmetropization. PLoS ONE 9, (2014).

2. Ellett, F., Pase, L., Hayman, J. W., Andrianopoulos, A. & Lieschke, G. J. mpeg1 promoter transgenes direct macrophage-lineage expression in zebrafish. Blood 117, e49–e56 (2011).

3. Eom, D. S. & Parichy, D. M. A macrophage relay for long distance signaling during post-embryonic tissue remodeling. Science 355, 1317–1320 (2017).

4. Collery, R. F., Volberding, P. J., Bostrom, J. R., Link, B. A. & Besharse, J. C. Loss of Zebrafish Mfrp Causes Nanophthalmia, Hyperopia, and Accumulation of Subretinal Macrophages. Invest. Ophthalmol. Vis. Sci. 57, 6805–6814 (2016).

5. Almoallem, B. et al. The majority of autosomal recessive nanophthalmos and posterior microphthalmia can be attributed to biallelic sequence and structural variants in MFRP and PRSS56. Sci. Rep. 10, (2020).

6. Bacci, G. M. et al. Novel mutations in MFRP and PRSS56 are associated with posterior microphthalmos. Ophthalmic Genet. 0, 1–8 (2020).

7. Crespí, J. et al. A Novel Mutation Confirms MFRP as the Gene Causing the Syndrome of Nanophthalmos-Renititis Pigmentosa-Foveoschisis-Optic Disk Drusen. Am. J. Ophthalmol. 146, 323–328.e1 (2008).

8. Guo, C. et al. Detection of Clinically Relevant Genetic Variants in Chinese Patients With Nanophthalmos by Trio-Based Whole-Genome Sequencing Study. Invest. Ophthalmol. Vis. Sci. 60, 2904–2913 (2019).

9. Katoh, M. Molecular Cloning and Characterization of MFRP, a Novel Gene Encoding a Membrane-Type Frizzled-Related Protein. Biochem. Biophys. Res. Commun. 282, 116–123 (2001).

10. Matsushita, I., Kondo, H. & Tawara, A. Novel compound heterozygous mutations in the MFRP gene in a Japanese patient with posterior microphthalmos. Jpn. J. Ophthalmol. 56, 396–400 (2012).

11. Sundin, O. H. et al. Extreme hyperopia is the result of null mutations in MFRP, which encodes a Frizzled-related protein. Proc. Natl. Acad. Sci. 102, 9553–9558 (2005).

12. Wasmann, R. A. et al. Novel membrane frizzled-related protein gene mutation as cause of posterior microphthalmia resulting in high hyperopia with macular folds. Acta Ophthalmol. (Copenh.) 92, 276–281 (2014).

13. Zenteno, J. C., Buentello-Volante, B., Quiroz-González, M. A. & Quiroz-Reyes, M. A. Compound heterozygosity for a novel and a recurrent MFRP gene mutation in a family with the nanophthalmos-retinitis pigmentosa complex. Mol. Vis. 15, 1794–1798 (2009).

14. Fogerty, J. & Besharse, J. C. 174delG Mutation in Mouse MFRP Causes Photoreceptor Degeneration and RPE Atrophy. Invest. Ophthalmol. Vis. Sci. 52, 7256–7266 (2011).

15. Won, J. et al. Membrane Frizzled Related Protein is necessary for the normal development and maintenance of photoreceptor outer segments. Vis. Neurosci. 25, 563–574 (2008).

16. Krebs, M. P., Hicks, W. & Nishina, P. M. Mfrp regulates ocular growth in mice and interacts with Prss56. Invest. Ophthalmol. Vis. Sci. 57, 3609–3609 (2016).

17. Velez, G. et al. Gene Therapy Restores Mfrp and Corrects Axial Eye Length. Sci. Rep. 7, (2017).

18. Fogerty, J. & Besharse, J. C. Subretinal infiltration of monocyte derived cells and complement misregulation in mice with AMD-like pathology. Adv. Exp. Med. Biol. 801, 355–363 (2014).

19. Zenteno, J. C., Buentello-Volante, B., Quiroz-González, M. A. & Quiroz-Reyes, M. A. Compound heterozygosity for a novel and a recurrent MFRP gene mutation in a family with the nanophthalmos-retinitis pigmentosa complex. Mol. Vis. 15, 1794–1798 (2009).

20. Curado, S., Stainier, D. Y. R. & Anderson, R. M. Nitroreductase-mediated cell/tissue ablation in zebrafish: a spatially and temporally controlled ablation method with applications in developmental and regeneration studies. Nat. Protoc. 3, 948–954 (2008).

21. Becker, T. & Becker, C. G. Regenerating descending axons preferentially reroute to the gray matter in the presence of a general macrophage/microglial reaction caudal to a spinal transection in adult zebrafish. J. Comp. Neurol. 433, 131–147 (2001).

22. Redd, M. J., Kelly, G., Dunn, G., Way, M. & Martin, P. Imaging macrophage chemotaxis in vivo: Studies of microtubule function in zebrafish wound inflammation. Cell Motil. 63, 415–422 (2006).

23. Ouyang, X. et al. The collagen metabolism affects the scleral mechanical properties in the different processes of scleral remodeling. Biomed. Pharmacother. 118, 109294 (2019).

24. Zhao, F. et al. Up-Regulation of Matrix Metalloproteinase-2 by Scleral Monocyte-Derived Macrophages Contributes to Myopia Development. Am. J. Pathol. 190, 1888–1908 (2020).

25. Nicol, C. S., Barrow, J. & Redmond, A. Flagyl (8823 R.P.) in the Treatment of Trichomoniasis. Sex. Transm. Infect. 36, 152–153 (1960).

26. Watt, L. & Jennison, R. F. Clinical Evaluation of Metronidazole. Br. Med. J. 2, 902–905 (1960).

27. Löfmark, S., Edlund, C. & Nord, C. E. Metronidazole Is Still the Drug of Choice for Treatment of Anaerobic Infections. Clin. Infect. Dis. 50, S16–S23 (2010).

28. Manickam, R., Oh, H. Y. P., Tan, C. K., Paramalingam, E. & Wahli, W. Metronidazole Causes Skeletal Muscle Atrophy and Modulates Muscle Chronometabolism. Int. J. Mol. Sci. 19, 2418 (2018).

29. Li, X. et al. Pineal Photoreceptor Cells Are Required for Maintaining the Circadian Rhythms of Behavioral Visual Sensitivity in Zebrafish. PLOS ONE 7, e40508 (2012).

30. Simões, F. C. et al. Macrophages directly contribute collagen to scar formation during zebrafish heart regeneration and mouse heart repair. Nat. Commun. 11, 1–17 (2020).

